# *In silico* molecular target prediction unveils mebendazole as a potent MAPK14 inhibitor

**DOI:** 10.1101/2020.05.18.101329

**Authors:** Jeremy Ariey-Bonnet, Kendall Carrasco, Marion Le Grand, Laurent Hoffer, Stéphane Betzi, Mickael Feracci, Philipp Tsvetkov, Francois Devred, Yves Collette, Xavier Morelli, Pedro Ballester, Eddy Pasquier

## Abstract

The concept of polypharmacology involves the interaction of drug molecules with multiple molecular targets. It provides a unique opportunity for the repurposing of already-approved drugs to target key factors involved in human diseases. Herein, we used an *in silico* target prediction algorithm to investigate the mechanism of action of mebendazole, an anti-helminthic drug, currently repurposed in the treatment of brain tumors. First, we confirmed that mebendazole decreased the viability of glioblastoma cells *in vitro*. Our *in silico* approach unveiled 21 putative molecular targets for mebendazole, including 12 proteins significantly up-regulated at the gene level in glioblastoma as compared to normal brain tissue. Validation experiments were performed on three major kinases involved in cancer biology: ABL1, MAPK1/ERK2 and MAPK14/p38α. Mebendazole could inhibit the activity of these kinases *in vitro* in a dose-dependent manner, with a high potency against MAPK14. Its direct binding to MAPK14 was further validated *in vitro* and inhibition of MAPK14 kinase activity was confirmed in live glioblastoma cells. Consistent with biophysical data, molecular modeling suggested that mebendazole was able to bind to the catalytic site of MAPK14. Finally, gene silencing demonstrated that MAPK14 is involved in glioblastoma tumor spheroid growth and response to mebendazole treatment. This study thus highlighted the role of MAPK14 in the anticancer mechanism of action of mebendazole and provides further rationale for the pharmacological targeting of MAPK14 in brain tumors. It also opens new avenues for the development of novel MAPK14/p38α inhibitors to treat human diseases.

**Significance Statement:** This study provides a framework to investigate drug polypharmacology by rapidly identifying novel molecular targets of already-approved drugs. It unveils a new mechanism involved in the anticancer activity of anti-helminthic drug, mebendazole, which is currently being repurposed for the treatment of brain tumors. By helping to decipher the mechanism(s) of action of repurposed drugs in their new indications, this approach could contribute to the development of safer and more effective therapeutic strategies in oncology and beyond.

## Introduction

The concept of polypharmacology, which involves the interaction of drug molecules with multiple targets, has emerged in recent years as a new paradigm in drug development [1, 2]. Polypharmacology can either limit or expand the medical indications of pharmacological agents. On the one hand, unintended drug-target interactions can cause toxic side effects, which can restrict and even prevent clinical use of a given drug. On the other hand, multi-targeting activity can open new therapeutic avenues for already-approved drugs – a concept called drug repurposing. An early estimation of the degree of drug polypharmacology based on a dataset containing 5,215 drug-target associations and 557 targets was an average of 6.3 molecular targets per approved drug [3]. Recently, Peon et *al*. updated this estimation to an average of 11.5 molecular targets per drug in cells owing to a more complete dataset comprising 8,535 drug-target associations and 1,427 targets [4]. This high degree of polypharmacology thus provides a unique opportunity for the repurposing of already-approved drugs to target key factors involved in human diseases.

One of the main limitations of drug development is its elevated cost. Thus, the rapid identification of novel therapeutic targets for already-approved drugs provides a route to reduce development cost by expanding their indications to human conditions in which those targets have been validated [5, 6]. Here, we applied a multi-pronged approach to unveil and validate new molecular targets for a well-known repurposed drug called mebendazole (MBZ). This anti-helminthic agent, which belongs to the benzimidazole class, has been shown to display potent anti-cancer properties in various models of human cancers [7-12] and thus appears as promising candidate for drug repurposing in oncology. Although the discovery of its therapeutic potential in brain tumors was fortuitous [11], it already resulted in 3 ongoing clinical trials in high-grade gliomas in both adult and pediatric patients (NCT01729260, NCT02644291 and NCT01837862). Several mechanisms of action have been proposed to explain the anticancer properties of MBZ. These include tumor angiogenesis inhibition [12, 13], targeting of critical pathways involved in cancer such as Hedgehog signaling [14] and stimulation of anticancer immune response [15, 16]. Most of these effects have been linked to the ability of MBZ to induce microtubule depolymerization in cancer cells [8, 11, 17, 18]. However, the affinity of MBZ for human tubulin is lower than that of helminthic tubulin [19]. Furthermore, the toxic side effects of MBZ are different and significantly milder than those of conventional microtubule-targeting chemotherapy agents, taxanes and *Vinca* alkaloids. This strongly suggests that additional, yet unknown mechanisms may be involved in the anticancer activity of MBZ, warranting further investigation.

Herein, we used *in silico* drug target prediction to identify novel putative molecular targets of MBZ in glioblastoma (GBM) cells. Through experimental validation of the predicted targets, we discovered that MBZ binds to MAPK14/p38α and inhibits its kinase activity *in vitro* and *in cellulo*. In accordance with biophysical characterization, molecular modeling studies predicted that MBZ was able to bind the catalytic site of MAPK14. Finally, gene silencing by RNA interference confirmed that MAPK14 plays a key role in the cytotoxic activity of MBZ against GBM cells and represents a promising therapeutic target in GBM.

## Results

### 1. Mebendazole exerts potent anti-proliferative effects against GBM cell lines in vitro

To investigate the anti-proliferative properties of benzimidazole agents *in vitro*, a range of human glioblastoma (GBM) cell lines were used (U87, U87vIII, T98G, U251). As shown in **Figure 1**, all tested compounds exerted dose-dependent anti-proliferative effects against all 4 GBM cell lines. Potency varied significantly between the different benzimidazoles, except in U251 cell line, which was highly sensitive to all 4 compounds. In all tested cell lines, the most active compound was mebendazole (MBZ) and the least potent was albendazole. MBZ displayed strong antiproliferative activity with IC_50_ values ranging from 288 +/- 3 nM for U251 cell line to 2.1 +/- 0.6 μM for the most resistant cell line, T98G.

**Figure 1.**
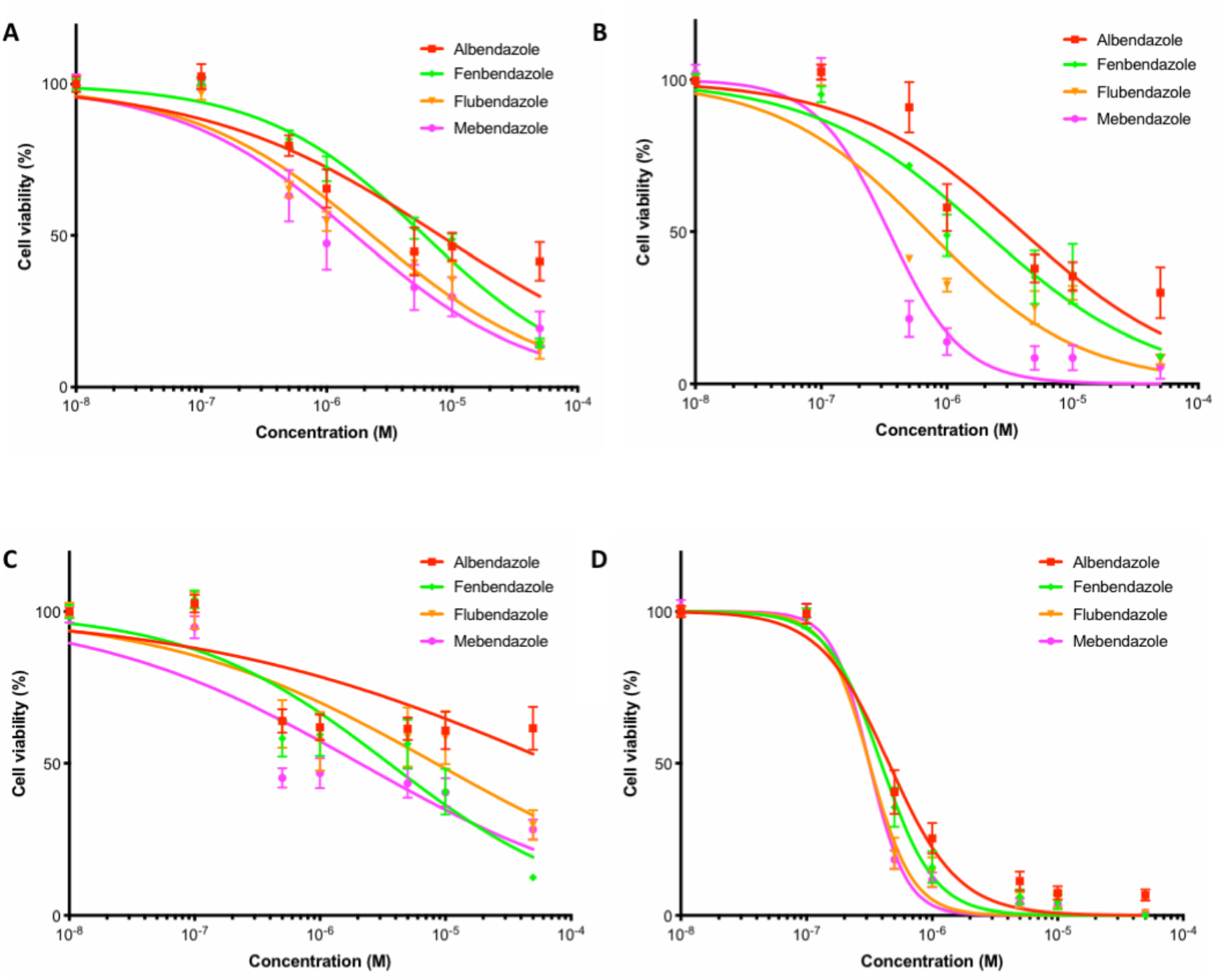
Impact of benzimidazole agents on glioblastoma cell viability *in vitro*. U87 (**A**), U87vIII (**B**), T98G (**C**) and U251 (**D**) GBM cells were incubated for 72h with increasing concentrations of benzimidazole agents. Cell viability was assessed by Alamar Blue assay and expressed as percentage of untreated cells. *Points*, mean of at least 4 independent experiments; *Bars*, sem.

### 2. In silico target prediction suggests multiple novel molecular targets for MBZ

To identify new therapeutic targets for MBZ, we used a data-driven predictive tool called MolTarPred [20]. Briefly, MolTarPred (http://moltarpred.marseille.inserm.fr/) is a freely-available web tool for predicting protein targets of small organic molecules. It is powered by a large knowledge base comprising 607,659 molecules and their known targets from the ChEMBL database [21]. MolTarPred returns the most similar target-annotated molecules to the user-supplied query molecule. Predicted targets are those known in similar molecules, thus potentially predicting up to 4,553 different protein targets [4]. Here, we used this *in silico* tool to unveil novel putative molecular targets for MBZ (**Figure 2**). We identified 4 other benzimidazole drugs, namely fenbendazole, flubendazole, albendazole and nocodazole, among the 10 molecules most similar to MBZ (out of 607,659 target-annotated molecules). MolTarPred predicted 21 human targets for this query molecule (**Table 1**). These include proteins involved in various biological processes, such as cell response to stress, signal transduction and cell cycle regulation. Interestingly, some of these proteins are major established therapeutic targets in oncology (*e.g*. TP53, MTOR/FRAP1, HIF1-α, VEGFR2/KDR and ABL1).

**Table 1.** List of putative mebendazole targets identified by *in silico* target prediction. *\*although KDR/VEGFR2 is a membrane receptor, its kinase activity is inhibited by benzimidazole agents* [Dakshanamurthy, 2012]

| Gene name | Full protein name | Main function |
| --- | --- | --- |
| **ABL1** | Tyrosine-protein kinase ABL | **Kinase** |
| **MAPK1** | MAP kinase ERK2 |  |
| **MAPK14** | MAP kinase p38 alpha |  |
| **FRAP1** | Serine/threonine-protein kinase mTOR |  |
| **TEK** | Tyrosine-protein kinase TIE-2 |  |
| **KDR\*** | Vascular endothelial growth factor receptor 2 |  |
| **LGR3** | Thyroid stimulating hormone receptor | **Receptor** |
| **ADORA2A** | Adenosine A2a receptor |  |
| **NPSR1** | Neuropeptide S receptor |  |
| **NPC1** | Niemann-Pick C1 protein | **Transporter** |
| **SLC6A3** | Dopamine transporter |  |
| **HIF1A** | Hypoxia-inducible factor 1 alpha | **Transcription factor** |
| **NFKB1** | Nuclear factor NF-kappa-B p105 subunit |  |
| **CREBBP** | CREB-binding protein | Transcription co-activator |
| **LMNA** | Prelamin-A/C | Nuclear membrane |
| **GMNN** | Geminin | DNA replication inhibitor |
| **BLM** | Bloom syndrome protein | DNA helicase |
| **TP53** | Cellular tumor antigen p53 | Tumor suppressor |
| **ALDH1A1** | Aldehyde dehydrogenase 1A1 | Enzyme |
| **THPO** | Thrombopoietin | Hormone |
| **HTT** | Huntingtin | vesicular transport |

**Figure 2.**
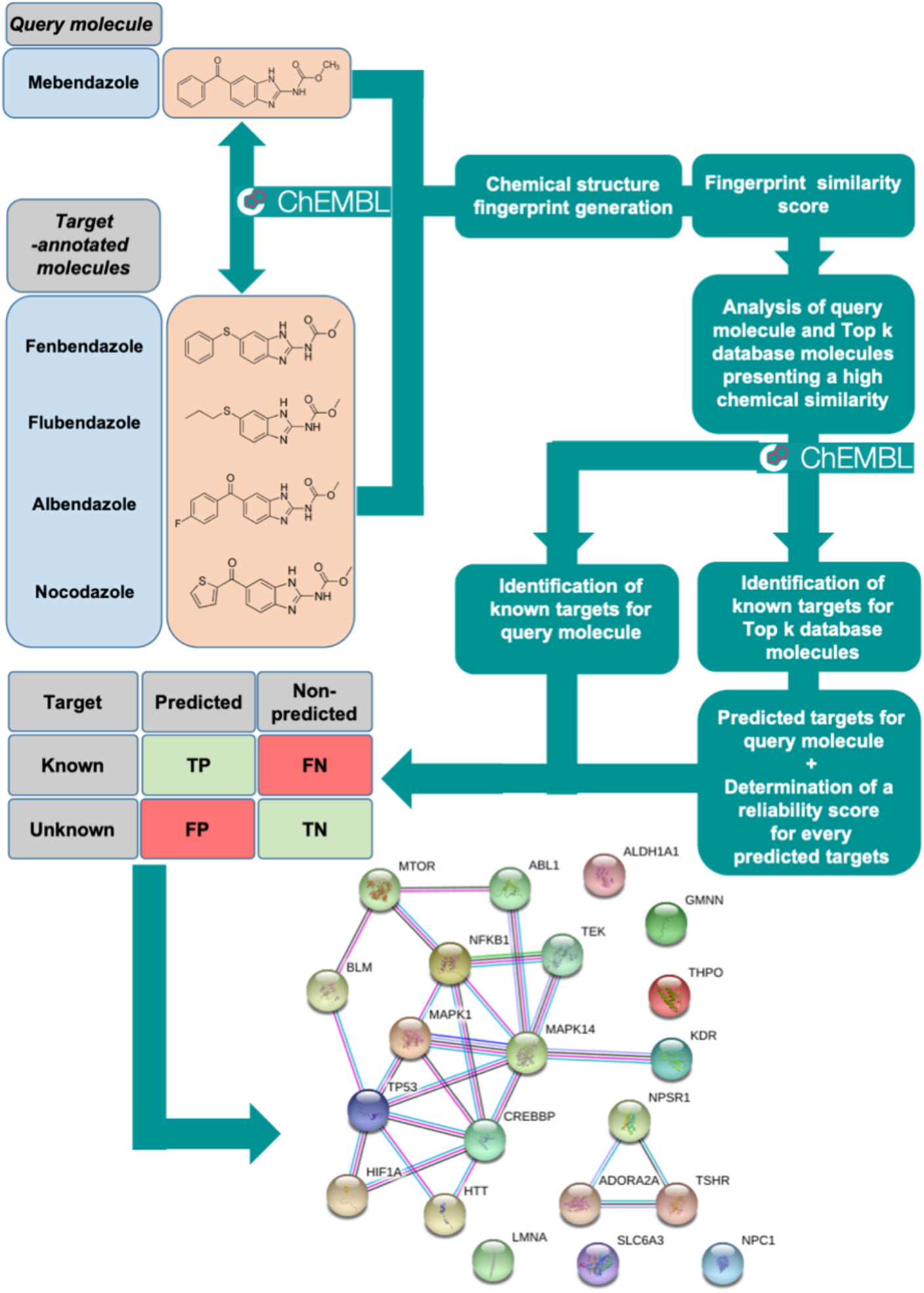
MolTarPred ligand-centric target prediction workflow. The most similar database molecules to the query molecule (mebendazole) are identified as previously detailed [Peon, 2019]. Known targets for these molecules are retrieved from the ChEMBL database. A reliability score for each query-target association prediction is calculated based on the proportion of the query’s top hits binding to the predicted target. The protein-protein interaction network of the 21 human proteins predicted as targets of MBZ is provided (string-db.org, Version 11.0).

### 3. MBZ inhibits ABL1, ERK2/MAPK1and MAPK14/p38α in vitro, with particularly high potency against MAPK14/p38α

To prioritize the putative molecular targets of MBZ, we performed transcriptomic analyses using freely available databases. We found that 12/21 putative MBZ targets were significantly up-regulated by at least 1.5 fold in GBM as compared to normal brain tissue (**Supp table 1**). Kinases have been amongst the most intensively pursued targets in oncology, leading to the approval of 46 kinase inhibitors for cancer treatment to date and over 150 kinase inhibitors currently in clinical trials [22, 23]. Interestingly, 1/3 of the putative MBZ targets up-regulated in GBM are kinases: ABL1, MAPK14/p38α, ERK2/MAPK1 and VEGFR2/KDR. Since MBZ has been previously shown to inhibit the kinase activity of the latter *in vitro* [24], we focused our validation experiments on the other 3 kinases (**Figure 3**). To experimentally validate the results of the *in silico* prediction, we performed functional assays to determine whether benzimidazoles were able to directly inhibit the activity of these kinases. We thus performed *in vitro* kinase assays on ABL1, MAPK14 and ERK2 kinases using a large range of concentrations (0,01 nM to 100 μM) of MBZ, albendazole, fenbendazole, flubendazole and nocodazole. As illustrated in **Figure 4**, benzimidazoles were able to inhibit the kinase activity of ABL1 and MAPK14 in a dose-dependent manner and to a lesser extent for ERK2. All tested compounds showed differential potency against the three kinases, with nocodozale being the most potent at inhibiting ABL1 activity (IC_50_ = 78 +/- 34 nM), MBZ at inhibiting MAPK14 activity (IC_50_ = 104 +/- 46 nM), and albendazole at inhibiting ERK2 activity (IC_50_ = 3.4 +/- 1.5 μM). The different IC_50_ values are summarized in **Table 2**. Collectively, these results confirm that MBZ inhibits all three tested predicted targets *in vitro*, with particularly high potency against MAPK14/p38α.

**Table 2.** IC_50_ values of benzimidazole agent in ABL1, ERK2 and MAPK14 *in vitro* kinase assays

| Drug | IC<sub>50</sub> value (μM) +/- SD |  |  |
| --- | --- | --- | --- |
|  | ABL1 | ERK2 | MAPK14 |
| Albendazole | 0.9 +/- 0.6 | 3.4 +/- 1.5 | > 50 |
| Fenbendazole | 0.19 +/- 0.07 | > 50 | 10.7 +/- 4.9 |
| Flubendazole | 0.11 +/- 0.05 | 15.9 +/- 8.0 | 5.5 +/- 2.7 |
| Mebendazole | 0.35 +/- 0.12 | 24.7 +/- 10.4 | 0.1 +/- 0.05 |
| Nocodazole | 0.08 +/- 0.03 | 26.9 +/- 16.2 | 10.8 +/- 5.6 |

**Figure 3.**
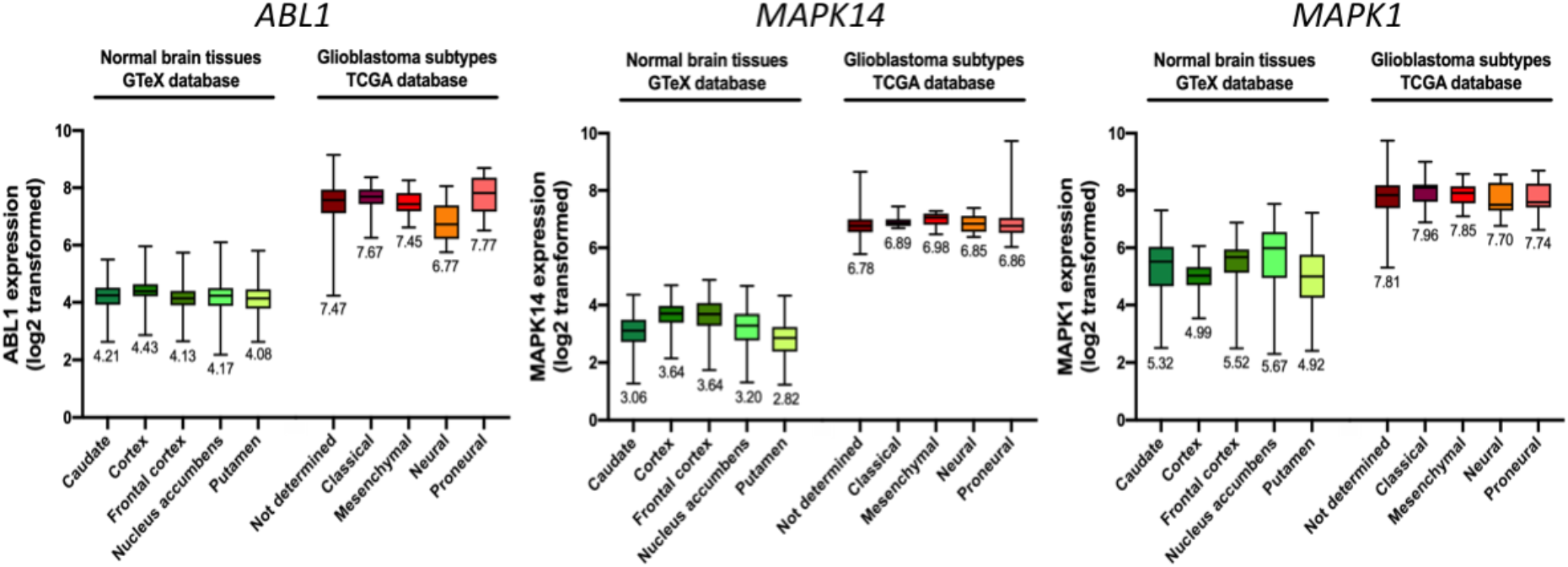
*ABL1, MAPK14* and *MAPK1* gene expression in glioblastoma and normal brain tissue. Normal brain tissue and GBM tissue gene expression values were obtained from the GTeX and TCGA databases. Boxplots representation showing relative Log2 transformed gene expression for *ABL1* (*lef*), *MAPK14* (*middle*) and *MAPK1* (*right*) in normal brain tissue (caudate (n=246), cortex (n=255), frontal cortex (n=209), nucleus accumbens (n=246) and putamen (n=205)) and GBM subtypes (not determined (n=455), classical (n=17), mesenchymal (n=27), neural (n=17) and proneural (n=24)) are shown. Average gene expression value is indicated below each boxplot.

**Figure 4.**
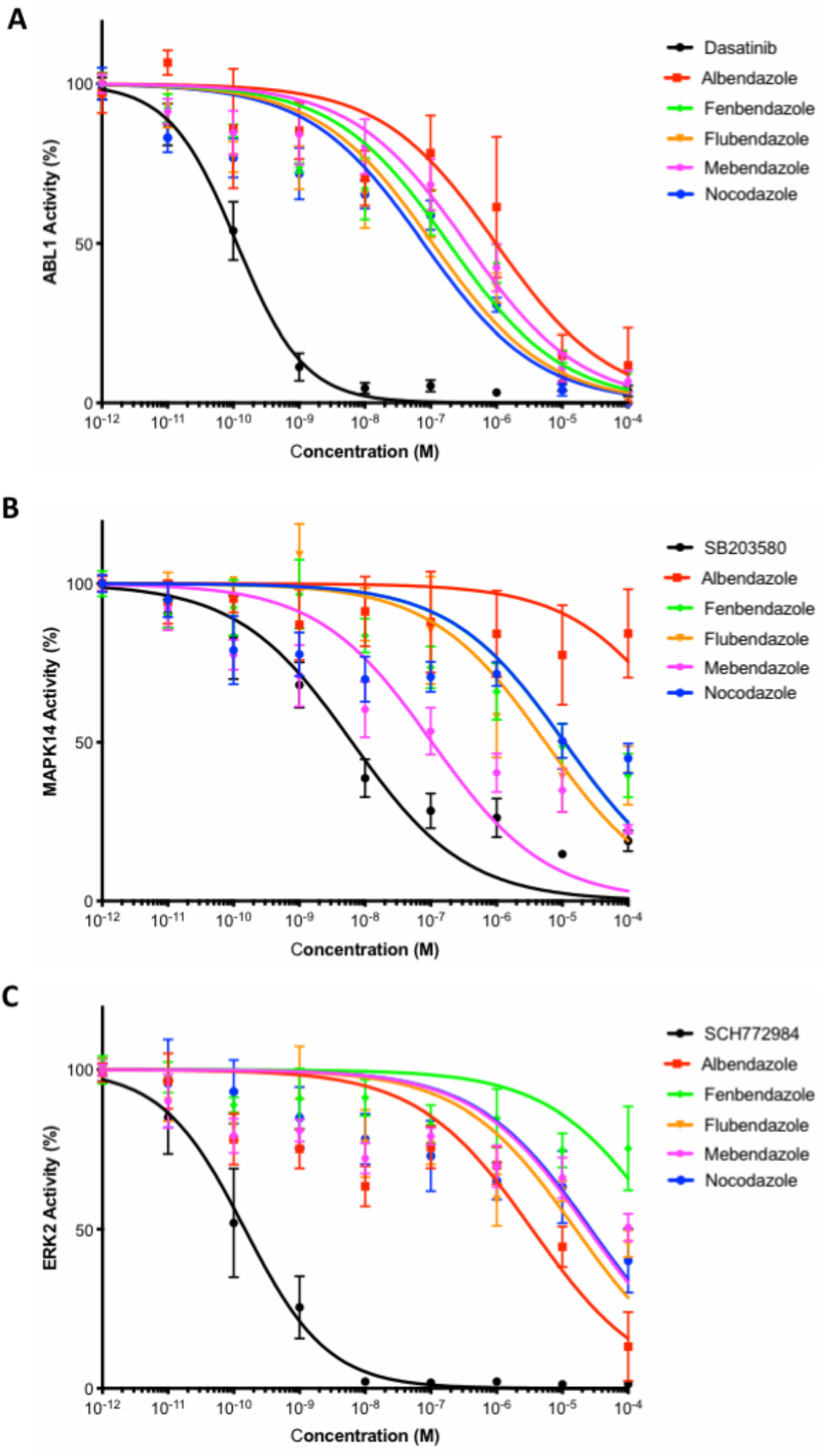
*In vitro* validation of protein kinase inhibition by benzimidazole agents. The concentration-dependent inhibition of ABL1 (**A**), MAPK14 (**B**), and ERK2 (**C**) by benzimidazole agents was determined by kinase assay as described in *Material and Methods*. Kinase activities in the absence of inhibitor were set to 100%, and remaining activities at different drug concentrations are expressed relative to this value. Dasatinib, SB203580 and SCH772984 were used as positive control to inhibit the kinase activity of ABL1, MAPK14 and ERK2, respectively. *Points*, mean of at least 4 independent experiments; *Bars*, sem.

### 4. MBZ interacts directly with MAPK14 in vitro and inhibits its kinase activity in cellulo

To characterize the interaction of MBZ with MAPK14 and perform orthogonal validation, we then employed a panel of established biophysical techniques, including Thermal Shift Assay (TSA), nanoscale Differential Scanning Fluorimetry (nanoDSF) and Isothermal Titration Calorimetry (ITC). As represented in **Figure 5A-B**, MAPK14 selective inhibitor, SB203580, was used as a positive control and induced an increase in MAPK14 thermostability at 50 μM (+12.7°C). MBZ was also able to increase the thermostability of MAPK14, which shifted from 44.3°C for the free form to 48°C in presence of MBZ at 50 μM (+3.7°C). Similarly, MBZ was able to increase the thermostability of ABL1 (+6.3°C) but to a lower extent than clinically-approved ABL1 inhibitors, imatinib and dasatinib (+12.3 and +18.8°C, respectively; **Supp Figure 1**). The direct binding of MBZ to MAPK14 was confirmed by nanoDSF, where increasing drug concentrations resulted in increased thermostability (**Figure 5C**). Furthermore, MBZ binding to MAPK14 was validated by ITC measurements, which allowed us to quantify the dissociation constant of the MBZ-MAPK14 complex (1.27 +/- 0.02 μM; **Figure 5D**). The thermodynamics parameters measured for MBZ exhibit a strong enthalpy component of −14 kcal.mol^−1^ compared to the −11.75 kcal.mol^−1^ measured for SB203580 (**Supp Figure 2**). Therefore, MBZ should engage the target with a favorable network of hydrogen bond interactions. SB203580 better Kd (0.18 +/- 0.01 μM; **Figure 5D**) is explained by a compensated favorable entropy component that could be partly explained by the compound rigidity. Finally, in order to demonstrate that MBZ was able to inhibit MAPK14 activity in live GBM cells, we conducted NanoBRET^™^ target engagement intracellular kinase assay in U87 cells transfected with MAPK14-NanoLuc^®^ Fusion Vector DNA. Here, we found that the BRET ratio decreased after MBZ treatment in a dose-dependent manner (IC_50_ = 4.1 +/- 1.1 μM). Taken together, these results show that MBZ is able to directly bind to MAPK14 and inhibit its kinase activity *in vitro* and *in cellulo*.

**Figure 5.**
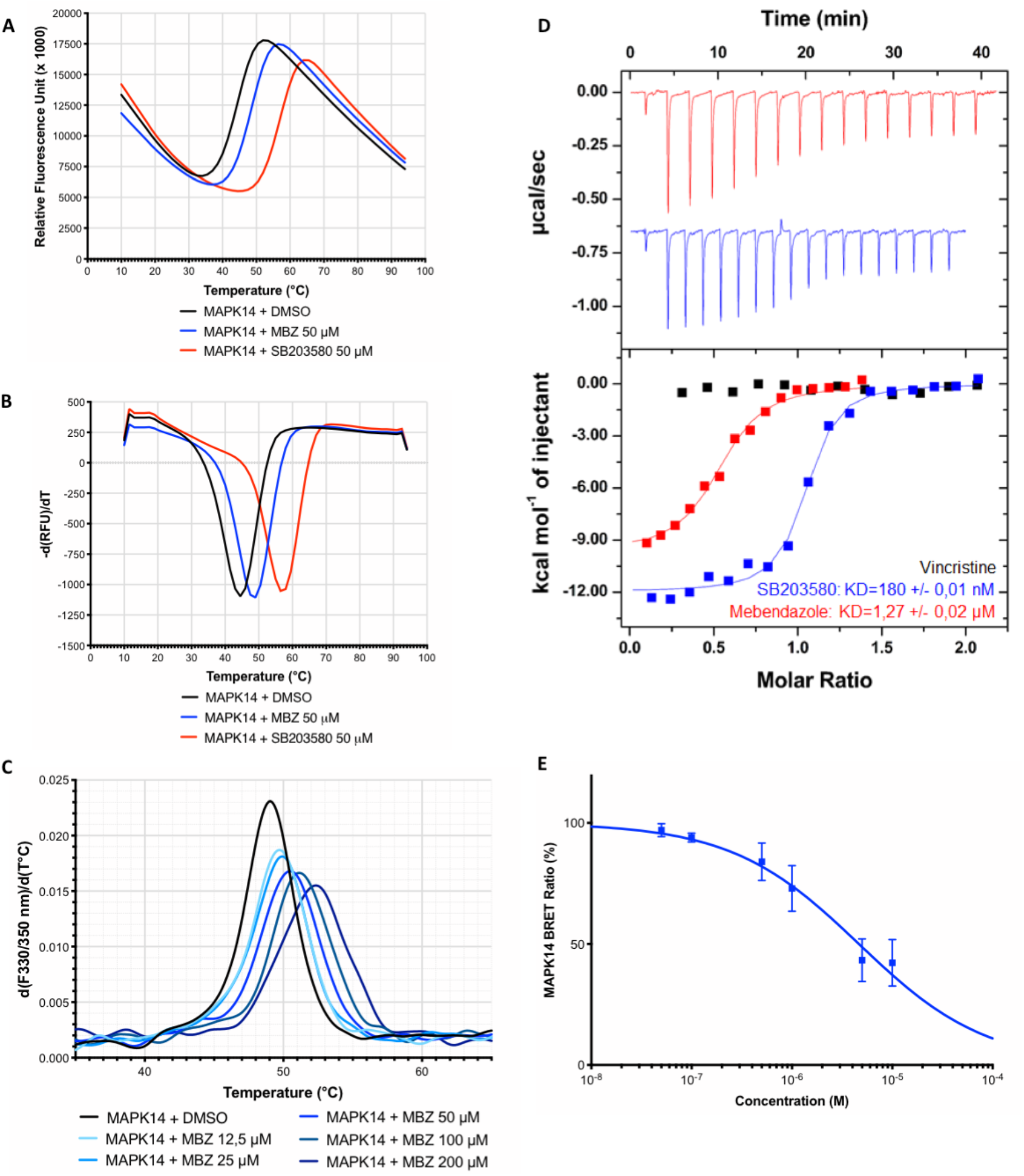
Biophysical characterization of MBZ binding to MAPK14 *in vitro* and *in cellulo* validation. Representative unfolding curves (**A**) and positive derivative [d(RFU)/dT] curves (**B**) of fluorescence-based TSA performed on 5 μM of MAPK14 alone (*black*) or incubated with 50 μM of MBZ (*blue*) or SB203580 (*red*) over a temperature range of 10–95°C. (**C**) First-derivative curves of nanoDSF thermal shift on MAPK14 alone (5 μM) or in presence of various concentrations of MBZ. (**D**) ITC measurements of the interaction between MBZ and MAPK14 protein using vincristine and SB203580 as negative and positive control, respectively. (**E**) nanoBRET target engagement assay with U87 cells transiently transfected with MAPK14-NanoLuc fusion vector incubated with increasing concentrations of MBZ for 1h. *Points*, mean of at least 4 independent experiments; *Bars*, sem.

### 5. Prediction of the binding mode of MBZ within MAPK14 using molecular modeling

More than 200 structures of MAPK14 (UniProt ID Q16539) are available from the Protein Data Bank (PDB) structural database. Kinases are known to be highly flexible proteins and large structural rearrangements can be observed after the superimposition of all structures in the same referential. Docking simulations were performed on several PDB structures. The 3FLY one resulted in a coherent predicted binding mode for MBZ and its analog compounds. The post-processed binding mode using SeeSAR is displayed in **Figure 6** using both 3D and 2D imageries. As anticipated from the measured enthalpy data by ITC, several hydrogen bond interactions are predicted by the model. More precisely, the carbamate-benzimidazole moiety is expected to make two hydrogen bonds with backbone atoms from M109, a crucial hot spot of the ATP binding pocket. Besides, this aromatic core is also superimposed with the potent pyrido-pyrimidin inhibitor from the 3FLY structure (IC_50_=4nM). The carbonyl group between both aromatic rings of MBZ is also predicted, as the 3FLY ligand, to make a water-mediated hydrogen bond with the sidechain from D168 (**Supp Figure 3A**). This last feature was not included in the water-free docking experiments but was highlighted by the SeeSAR postprocessing where crystallographic waters from the reference PDB structure can be switched on/off. Finally, the terminal phenyl ring of MBZ is predicted to be superimposed with the difluorophenyl moiety from the 3FLY pyrido-pyrimidin inhibitor, deeply buried within a small and well-defined hydrophobic pocket delimited by V38, A51, K53, L75, L86, L104 residues. The molecular modeling study was also able to explain the lower affinity of albendazole for MAPK14 from its predicted binding mode. The single difference between albendazole and MBZ ligands is their terminal end: a rigid aromatic ring for MBZ and a flexible aliphatic chain for albendazole. Therefore, albendazole is predicted to make less Van der Waals interactions between its hydrophobic sidechain within the hydrophobic pocket (**Supp Figure 3B**). Besides, albendazole lacks the carbonyl spacer that is predicted to be involved in a water-mediated hydrogen bond with the protein. Finally, this ligand also exhibits more flexibility that is expected to decrease the affinity from an entropic point of view.

**Figure 6.**
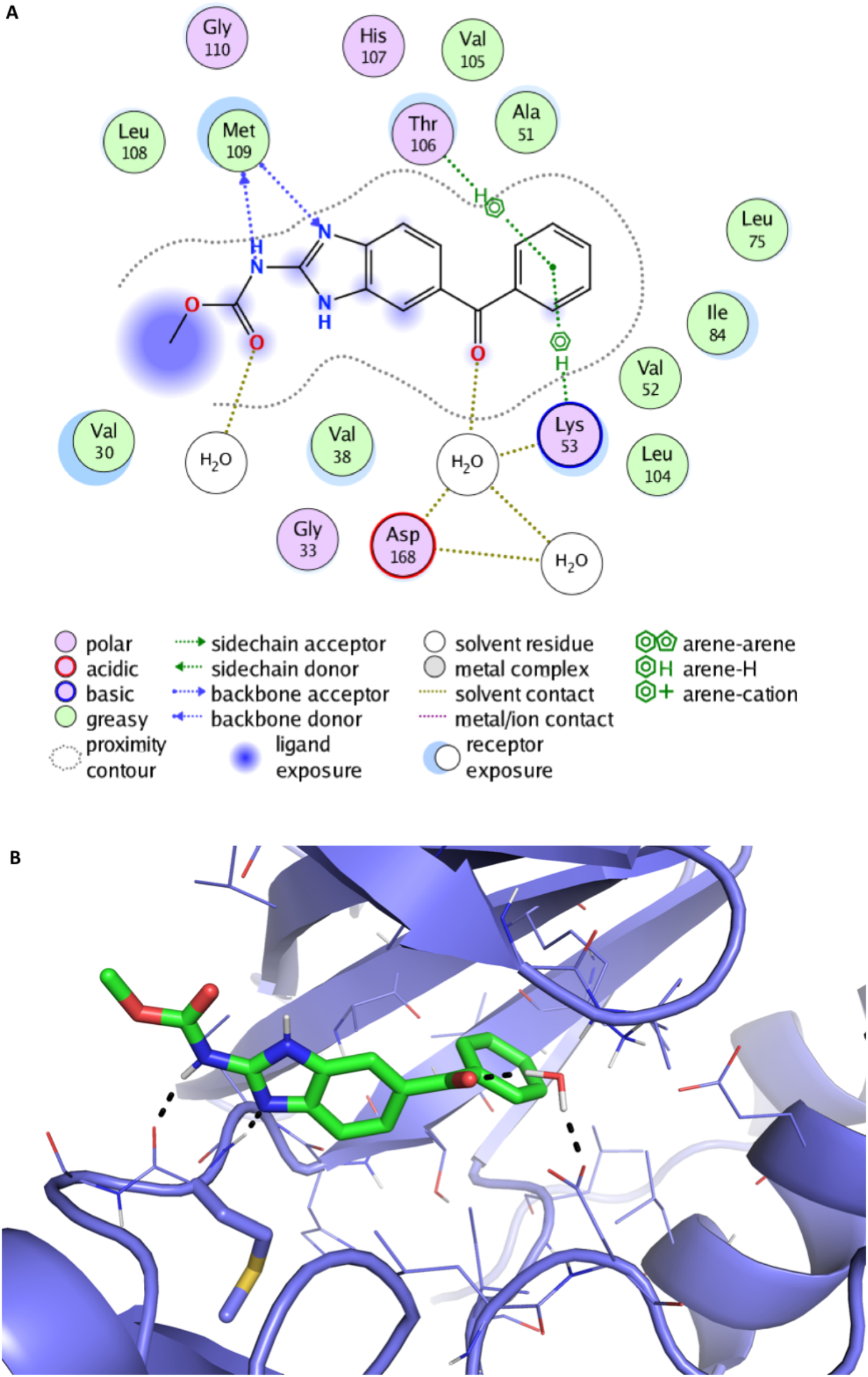
Proposed binding mode of mebendazole to MAPK14. The binding mode of MBZ to MAPK14 protein is depicted as 2D (**A**) and 3D diagrams (**B**), which were generated using the “Ligand interaction” feature from MOE and Pymol software, respectively. The benzimidazole core is predicted to make hydrogen-bond interaction with the hot spot M109 and the other aromatic moiety is deeply buried in a hydrophobic pocket of the binding site. MBZ and the sidechain from M109 are displayed in sticks and surrounding protein residues are depicted in lines. The backbone from the protein is shown as cartoon representation.

### 6. MAPK14 is a therapeutic target in GBM

The cytotoxic activity of benzimidazoles significantly correlated with their ability to inhibit MAPK14 kinase activity *in vitro* in 2 out 4 tested GBM cell lines and a similar trend was observed in the other 2 cell lines (**Supp Table 2**). Interestingly, this was not observed with ABL1 and ERK2 kinase activity. We therefore performed *MAPK14* gene silencing in GBM cells to confirm its role as a key molecular target of MBZ. *MAPK14* gene expression was knocked down in dsRed-expressing U87 cells by transfection with 3 different siRNA sequences (**Supp Figure 4**). This resulted in a significant decrease in tumor spheroid growth *in vitro* (**Figure 7A**). While negative control siRNA-transfected spheroids grew by 595 % in 7 days, they only grew by 260 - 460 % when transfected with MAPK14-targeting siRNA (p<0.05). Similarly, spheroid doubling time increased from 75h following transfection with negative control siRNA to 172, 149 and 100h following transfection with MAPK14 siRNA sequences #1, #2 and #3, respectively. Consistent with the role of MAPK14 as a critical molecular target of MBZ, *MAPK14* gene silencing significantly decreased the sensitivity of GBM cells to the drug (**Figure 7B**). The IC_50_ value of MBZ after 192h incubation increased from 5 and 6.6 μM in Mock and control siRNA-transfected cells, respectively to 17.6 μM in cells transfected with MAPK14-targeting siRNA sequence #3 and > 50 μM in cells transfected with the other 2 siRNA sequences. These results confirmed the role of MAPK14 in GBM cell response to MBZ.

**Figure 7.**
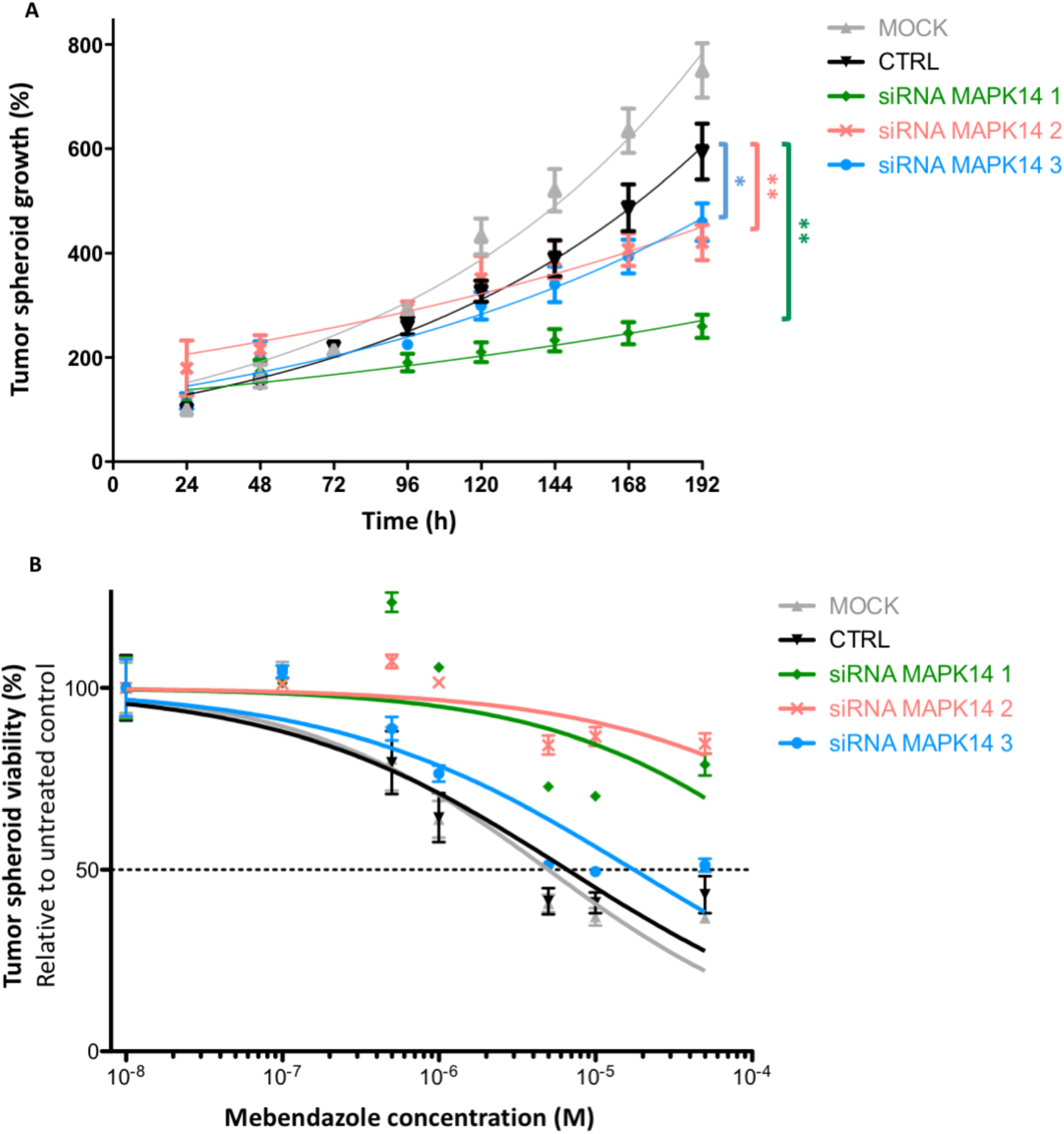
Functional validation of MAPK14 as a key molecular target of mebendazole in glioblastoma cells. (**A**) Growth curves of 3D tumor spheroids formed by dsRed-expressing U87 cells either untransfected (mock) or transfected with negative control or MAPK14-targeting siRNA. Spheroid growth was assessed by daily fluorescence measurements at 575/620 nm. (**B**) Dose-response curve of dsRed-expressing U87 cells, either untransfected (mock) or transfected with negative control or MAPK14-targeting siRNA, incubated with increasing concentrations of MBZ for 192h. *Points*, mean of at least 4 independent experiments; *Bars*, sem.

## Discussion

Drug repurposing, which consists in using already-approved drugs in new medical indications, has become an attractive therapeutic strategy in oncology. There are however two major hurdles to overcome for successful drug repurposing strategies: 1) identifying the right drug for the right situation (*i.e*. disease type, patient population, clinical setting…), and 2) deciphering the new mechanism(s) of action of repurposed drugs in the context of cancer. In this study, we established an innovative framework to rapidly identify and validate the molecular target(s) of MBZ, an anti-helminthic agent, currently repurposed for the treatment of GBM. Our results revealed for the first time a crucial role of MAPK14 in its mechanism of action.

The conventional approaches to drug development in oncology take on average 12 to 15 years and the median cost of anticancer drugs at the time of approval went from less than $100 per month in the 1990’s to over $10,000 per month currently, threatening national healthcare systems worldwide [5, 25]. In this context, drug repurposing approach holds great promise to develop safe, effective and inexpensive therapeutic options that can be made readily available to cancer patients regardless of their socio-economic status [5, 26]. Over the last decade, MBZ has been recognized as an attractive candidate for drug repurposing in various models of human cancers [7-12]. Its anti-cancer properties were notably demonstrated in gliomas [11, 14, 27] and there is currently 3 ongoing clinical trials in high-grade gliomas in both adult and pediatric patients (NCT01729260, NCT02644291 and NCT01837862). Furthermore, a recent phase I trial demonstrated that administration of MBZ concomitantly with radiotherapy and temozolomide chemotherapy was safe in high-grade glioma patients [28]

Although multiple studies have highlighted different mechanisms of action of MBZ in cancer cells [12-16], including its effects on the microtubule network [8, 11, 14, 17, 18], its precise molecular target(s) in tumor cells remained to be ascertained. Herein, we used *in silico* target prediction to unveil novel therapeutic targets for MBZ in GBM. We foresaw 21 putative targets, of which four had been previously shown to be modulated by benzimidazole agents *in vitro*: ABL1, VEGFR2, mTOR and aldehyde dehydrogenase [19, 24, 29, 30]. Our methodology thus identified 17 novel putative targets of MBZ. We also observed that 12 of the 21 predicted targets are significantly overexpressed in GBM as compared to normal brain tissue, which may explain why MBZ is particularly effective against brain tumors [11, 13, 14, 17, 31].

We focused our validation experiments on three major kinases involved in cancer: the tyrosine kinase ABL1, the Mitogen-Activated Protein Kinase (MAPK) member ERK2 and one of the four p38 MAPKs, MAPK14. All of these proteins have already been shown to play a key role in the pathophysiology or drug resistance of GBM. Indeed, while ABL1 was found to be involved in DNA repair upon irradiation, thus contributing to radioresistance in GBM cells [32], a crucial role in linking RNA processing with signal transduction of ERK2 has recently been unveiled [33]. Moreover, activation of the MAPK14 signaling pathway has been previously shown to correlate with poor prognosis in GBM patients, increased tumor invasiveness and aggressive phenotype [34-38]. Here, *in vitro* kinase assays confirmed that MBZ could directly inhibit the kinase activity of these proteins, thus confirming their status of direct molecular targets of MBZ. Moreover, we found that all 3 kinases are over-expressed in GBM as compared to normal brain, thus providing an opportunity for targeted strategy in this deadly form of brain tumor.

MAPK14 is one of the four p38 MAPKs that play an important role in the cascades of cellular responses evoked by extracellular stimuli such as proinflammatory cytokines or physical stress leading to direct activation of transcription factors [39]. For many years, p38 MAPK kinases have been considered as attractive targets for chronic inflammatory disease therapy. p38 MAPK inhibitors have taken a leap forward through the development of many compounds for different pathologies, including cancers [39, 40]. Here we have discovered that MBZ is particularly potent at inhibiting MAPK14 kinase activity *in vitro*. Molecular modeling studies also revealed that the carbamate-benzimidazole moiety of MBZ engages with the critical M109 hot spot of the catalytic site of MAPK14. Interestingly, it has been demonstrated that the binding mode of selective MAPK14 inhibitors is characterized by a compound-induced peptide flip between M109 and G110 [41]. Besides, some MBZ-urea inhibitors are known to bind the hinge region from other kinases such as VEGFR2 [42], and one reference structure is publicly available from the PDB database (PDB ID 2OH4). Interestingly, the carbamate-benzimidazole moiety adopts the same interaction pattern with the homologous hot spot from VEGFR2 protein. This structural data strengthens the reliability of the predicted binding mode of the MBZ series within the MAPK14 binding site.

The MAPK14/p38α signaling pathway has been classically considered a tumor suppressor. However, several studies have also demonstrated the pro-tumorigenic activities of MAPK14 which facilitates the survival and proliferation of tumor cells [39]. In our study, the use of RNA interference revealed that MAPK14 expression is crucial to GBM tumor growth in a spheroid model, confirming the therapeutic potential of targeting MAPK14 in GBM. Consistently, MAPK14 selective inhibitor, LY2228820 (Ralimetinib), has been shown to produce significant tumor growth delay in multiple cancer models, including GBM [43, 44] and has now entered clinical trials [45]. Moreover, recent evidence suggest that MAPK14 is also involved in drug resistance. For instance, response to cisplatin can be enhanced by MAPK14 inhibition, resulting in ROS-dependent upregulation of the JNK pathway in colon and breast cancer cells [46]. Similarly, MAPK14 confers resistance to irinotecan in TP53-defective colon cancer cells by inducing pro-survival autophagy [47]. In line with this, MAPK14 was found to play a critical role in regulating response to temozolomide treatment in GBM [48, 49]. These studies along with the results presented here strongly suggest that targeting MAPK14 with MBZ or other pharmacological inhibitors represents a promising strategy to enhance chemotherapy efficacy in cancer, including temozolomide efficacy against GBM.

In conclusion, we have found consistent evidence that further supports the use of MBZ as a promising repurposed drug in various tumor types, and especially in brain cancers. We have discovered for the first time that MBZ is a potent inhibitor of MAPK14, which would directly contribute to its anticancer properties in GBM. Our results could thus open new therapeutic avenues for the development of MAPK14 inhibitor combination therapies in GBM and other human diseases. More broadly, we have established a framework for the rapid identification and functional validation of novel molecular targets that could be applied to other repurposed drugs, and therefore enable the development of innovative therapeutic strategies for unmet medical needs.

## Materials and Methods

### Cell culture

U87, U87vIII, T98G and U251 are glioblastoma cell lines. They were grown in Dulbecco’s Modified Eagle Medium (ThermoFisher Scientific) containing 10% Fetal Calf Serum (FCS) and 1% pyruvate and 1% penicillin-streptomycin. They were routinely maintained in culture on 0.1% gelatin-coated flasks at 37°C and 5% CO_2_. Both cell lines were regularly screened and are free from mycoplasma contamination.

### Cell viability assay

Cell viability assays were performed as previously described [50]. Briefly, cells were seeded at 4,500 cells/well in 96-well plates. After 24h, cells were treated with a range of concentrations of benzimidazole agents and after 72h drug incubation, metabolic activity was detected by addition of Alamar blue and spectrophotometric analysis using a PHERAstar plate reader (BMG labtech). Cell viability was determined and expressed as a percentage of untreated control cells. The determination of IC_50_ values was performed by using the following equation: Y=100/(1+((X/IC_50_)^Hillslope)).

### Gene expression analysis on patient samples

Gene expression analysis was conducted using the R2 microarray analysis and visualization platform (http://r2.amc.nl). RNA-seq data were extracted from two independent cohorts providing open access to data acquired from various forms of cancer: the Cancer Genome Atlas (TCGA) database and from normal tissues: the Genotype-Tissue Expression (GTeX) database. GBM TCGA dataset was used and partitioned in five subtypes according to the data available: classical (n=17), mesenchymal (n=27), neural (n=17), proneural (n=24) and not determined (n=455). We used GTeX normal brain tissue data from the following subgroups: caudate (n=246), cortex (n=255), frontal cortex (n=209), nucleus accumbens (n=246) and ptamen (n=205). Median values were recorded using log2 transformation gene expression. Statistical analyses using ANOVA were performed to compare GBM subtypes gene expression to normal brain tissue gene expression. Boxplots were generated using GraphPad Prism 8.4.1.

### Kinase assay

MAPK14, ERK2 and ABL1 kinase assay were purchased from Promega. Enzyme, substrate, ATP and inhibitors were diluted in Kinase Buffer as per the manufacturer’s instructions. Kinase reaction was performed in 384-well plate in a final volume of 5 μL. Reaction was initiated using 1 μL of inhibitor for each concentration (1% DMSO), 1 μL of enzyme and 3 μL of substrate/ATP mix (60min, RT). Five μL of ADP-GloTM Reagent were used to stop kinase reaction by ATP depletion (40min, RT). Then, ADP formed by kinase reaction was detected by adding 10 μL of Kinase Detection Reagent (30min, RT). Luminescence was recorded using a PHERAstar plate reader. MAPK14, ERK2, and ABL1 kinases were used at optimized concentrations of 4, 3 and 1 ng/well, respectively. For the three proteins, ATP was used at 5 μM and DTT at 50 μM. The substrates of MAPK14, ABL1, and ERK2 were used at 0.2, 0.2 and 0.1 μg/μL.

### NanoBRET target engagement assay

U87 cells were transfected with MAPK14-NanoLuc fusion vector DNA (Promega) using Lipofectamine^^™^^ RNAI Max and following the manufacturer’s instructions. 17,000 cells/well were dispensed into a 96-well NBS plates and prepared with 0.1 μM of NanoBRET^™^ Tracer K-4 reagent. Cells were treated with MBZ at concentrations ranging from 0.05 to 10 μM and incubated at 37°C, 5% CO_2_ for 2h. A separate set of samples without tracer was prepared for background correction. Plate was equilibrated at RT during 15min. Complete substrate plus inhibitor solution in assay medium (Opti-MEMRI reduced serum medium, no phenol red) was prepared just before measuring BRET signal. 50 μL of 3X complete substrate plus inhibitor solution was added to each well of the 96-well plate, and incubated for 2-3min at RT. Donor emission wavelength (460 nm) and acceptor emission wavelength (610 nm) were measured using a PHERAstar plate reader.

### Protein Expression and Purification

For MAPK14 expression, we used the pET28a vector kindly provided by Drs Qi and Huang of the National Institute of Biological Sciences in China [51]. The expression and purification of wild type MAPK14 were carried out as previously reported [Bukhtiyarova, 2004]. For the Isothermal Titration Calorimetry (ITC) and the Thermal Shift Assay (TSA) the His tag was conserved, and the protein was concentrated to 17 mg/mL in 25mM Tris HCL pH7.4, 150mM NaCl, 5% glycerol, 10mM MgCl2 and 5mM DTT. The samples were immediately flash frozen in liquid nitrogen and stored at −80°C. A pET28a vector was also used to perform the expression of the kinase domain of human c-ABL (ABL1). The protein was expressed in E. coli strain BL21 (DE3) STAR in TB media with 50 μg/mL kanamycin and 34 μg/mL Chloramphenicol at 17°C overnight after induction with 0.2 mM of IPTG. The bacteria were disrupted by sonication on ice for 3min in lysis buffer (50mM Tris–HCl pH8, 500mM NaCl, glycerol 5%, 10mM imidazole and 0.1% BRIJ35) with EDTA-free protease inhibitor cocktail (Roche). The protein from the soluble fraction was loaded onto a HisTrap Ni-Nta column, washed with 5 column volumes of lysis buffer containing 10 mM imidazole, 5 volumes of 40 mM imidazole buffer and eluted with 5 volumes of a gradient imidazole buffer from 40 to 500 mM imidazole. The resulting protein was concentrated and purified by size-exclusion chromatography using Superdex 75 (GE Healthcare) with 10mM Tris pH8.0, 50mM NaCl and 1mM DTT.

### Thermal-Shift Assay (TSA)

TSA experiments were performed in triplicate in 384-well PCR plates (Bio-Rad). The reagents (compound, protein, and fluorophore) were dispensed using an Echo550 acoustic dispenser (Labcyte): 100 nL of compound (from a 100% DMSO stock at 10 mM) for a final concentration of 50 μM (0.5% final DMSO); 200 nL of Thermal Shift Dye (Thermo Fisher Scientific) diluted to a final concentration of 0.1%; 300 nL of MAPK14 (42 μM stock) for a final concentration of 5 μM. The final assay volume was completed to 19.5 μL with assay buffer (10 mM HEPES, pH 7.5, 500 mM NaCl) using a Multidrop Combi (Thermo Fisher Scientific). For TSA experiments on ABL1, the kinase domain of human c-ABL (ABL1) was mixed at a final concentration of 4μM with the inhibitor (200 μM final concentration / 2% DMSO) in a final volume of 20μL of the assay buffer (25mM HEPES pH7.5 / 150mM NaCl / 1mM DTT / 1% glycerol). For both assays, the Thermal Shift Dye was added at the end diluted to a final concentration of 0.1%. The plates were sealed with optical film (Ampliseal, Greiner) and centrifuged at 1000 rpm for 1min at 4°C. The thermal-melting experiments were carried using a CFX384 RTPCR (Bio-Rad). The plates were first equilibrated at 25°C for 1min; then heated using a 0.5°C steps ramp from 25 to 95°C using 25s equilibration. Raw fluorescence was measured, and the melting temperatures (Tm) were calculated using CFX Manager 3.1 (Bio-Rad).

### Isothermal-Titration Calorimetry (ITC)

ITC was used to determine the thermodynamics parameters of the binding between MAPK14 and the selected compounds. Titrations were carried out on a MicroCal ITC200 microcalorimeter (GE Healthcare). Each experiment was designed as normal (protein in the syringe and ligand in the cell) or reverse (protein in the syringe and ligand in the cell) titrations experiments using 13 to 17 injections at 15 or 25°C. Raw data were scaled after setting the titration-saturation-heat value to zero. Integrated raw ITC data were fitted to a one-site nonlinear least-squares-fit model using the MicroCal Origin plugin as implemented in Origin 9.1 (Origin Lab). Finally, ΔG and TΔS values were calculated from the fitted ΔH and KD values using the equations ΔG=-RTlnKD andΔG=ΔH-TΔS. Each experiment was performed as triplicates and data are presented as the mean ± s.d.

### Nano Differential Scanning fluorimetry assay

The thermostability of MAPK14 was measured in the presence of different concentrations of MBZ in 50 mM HEPES buffer in the presence of 200 mM NaCl and 2% DMSO at pH 7.5 using a label-free fluorimetric analysis with a Prometheus NT.Plex instrument (NanoTemper Technologies), as described previously [52]. NanoDSF grade capillaries were filled with a 10 μM solution of interest. The concentration of MBZ varied from 0.78 to 200 μM while MAPK14 concentration was fixed at 5 μM. Capillaries were loaded into the Prometheus NT.Plex and heated from 25 to 70°C with a 1 K/min heating rate at low detector sensitivity with an excitation power of 10%. Unfolding transition points (Tm) were determined from the first derivative of the changes in ratio between the emission wavelengths of tryptophan fluorescence at 330nm and 350nm, which were automatically identified by the Prometheus NT.Plex control software.

### Molecular modeling

The PDB files (1A9U and 3FLY) were prepared using MOE version 2016 (http://chemcomp.com). The binding site was defined as all residues with at least one atom within 10 Å radius from the crucial M109 known hot spot. The studied compounds, including co-crystalized ligands and MBZ-like inhibitors, were also prepared using MOE in order to generate random 3D conformers and compute required partial charges. PLANTS was used as the docking engine with its “Chemplp” scoring function to generate and evaluate the poses [53]. Its docking algorithm is based on a class of stochastic optimization algorithms called Ant Colony Optimization. This kind of algorithm, which mimics the behavior of ants finding the shortest path between food and their nest, can be used to efficiently sample the conformational space for docking purpose. Neither geometric nor pharmacophore constraints were added in simulations. Redocking experiments of reference ligands (SB203580 and pyrido-pyrimidin inhibitor) into their respective structures (1A9U and 3FLY) were beforehand performed to validate the docking settings and protocol. This control study is used to determine the best docking parameters for the target [54]. Both ligands were successfully redocked into their binding site with a RMSD value of less than 1Å between crystallized conformation and predicted pose. All poses from docking simulations were subjected to visual analysis using Pymol (http://pymol.org) and the binding mode of interest was selected accordingly. The latter should 1) contain polar interactions with crucial hot spot M109, 2) highlight shared binding mode between all active MBZ analogs, 3) be compatible with known SAR data, and 4) be reasonable in terms of docking score and explicit interactions with the binding site. The most promising poses were subjected to post-processing with the commercial SeeSAR software (http://biosolveit.de). Briefly, this computer-aided design tool, which was developed for modelers and chemists, relies on the HYDE method [55] to evaluate affinity contributions to the binding for each ligand atom. SeeSAR can then be used to 1) optimize docking poses within the binding site, 2) identify optimal and sub-optimal moieties from ligands, 3) analyze SAR data from 3D complex point of view, and 4) design and evaluate new analog compounds in order to optimize the series. In this work, SeeSAR was initially used to identify potential issues in prioritized poses, mainly desolvation penalties, low-quality hydrogen-bonds and sub-optimal torsions. Ultimately, SeeSAR was used to estimate the affinity of each compound from the MBZ series from their respective post-processed binding mode.

### Functional validation of MAPK14

DsRed-expressing U87 cells were seeded in T25 flask cell culture and transfected with 1 ml of Opti-MEM medium containing 1% of lipofectamine RNAiMax and 5nM of siRNA. Three different siRNA sequences targeting MAPK14 were used (Silencer^®^ Select s3585, s3586 and s3587; ThermoFisher Scientific) as well as a non-targeting negative control siRNA with no significant sequence similarity to mouse, rat or human gene sequences (Silencer^®^ Select AM4635). Two days later, cells were seeded in 96-well U bottom and low-binding plates in DMEM medium containing methylcellulose at 0.6 g/L and spheroids were treated with an increasing concentration of MBZ 24h after seeding. Spheroid viability was then evaluated daily for one week by measuring the dsRed fluorescence ratio (575nm: excitation wavelength / 620nm: emission wavelength) with a PHERAstar plate reader.

To evaluate the level of gene knock-down, cells were harvested 48h after transfection and total RNA was extracted using RNeasy kit (Qiagen), following the manufacturer’s instructions. Reverse transcription was performed with Onescript^®^ cDNA synthesis kit (Abm) and qRT-PCR was performed using SsoAdvanced Universal SYBR^®^ Green Supermix (Bio-Rad) and a CFX96^^™^^ Real-Time System device (Bio-Rad). Gene expression levels were determined using the ΔΔ*C*t method, normalized to the *YWHAZ* control gene. The following predesigned KiCqStart SYBR^®^ Green primers (Merck) were used: *MAPK14* (forward: 5’-AGATTCTGGATTTTGGACTG; reverse: 5’-CCACTGACCAAATATCAACTG) and *YWHAZ* (forward: 5’-AACTTGACATTGTGGACATC; reverse: 5’-AAAACTATTTGTGGGACAGC).

## Acknowledgments

This work was supported by a grant from the not-for-profit organization “Eva pour la Vie” and by a “Pepiniere d’Excellence” grant from the A*MIDEX Foundation of Aix Marseille University, funded by socio-economic partners, both attributed to EP.

---

**Supplementary Figure 1.**
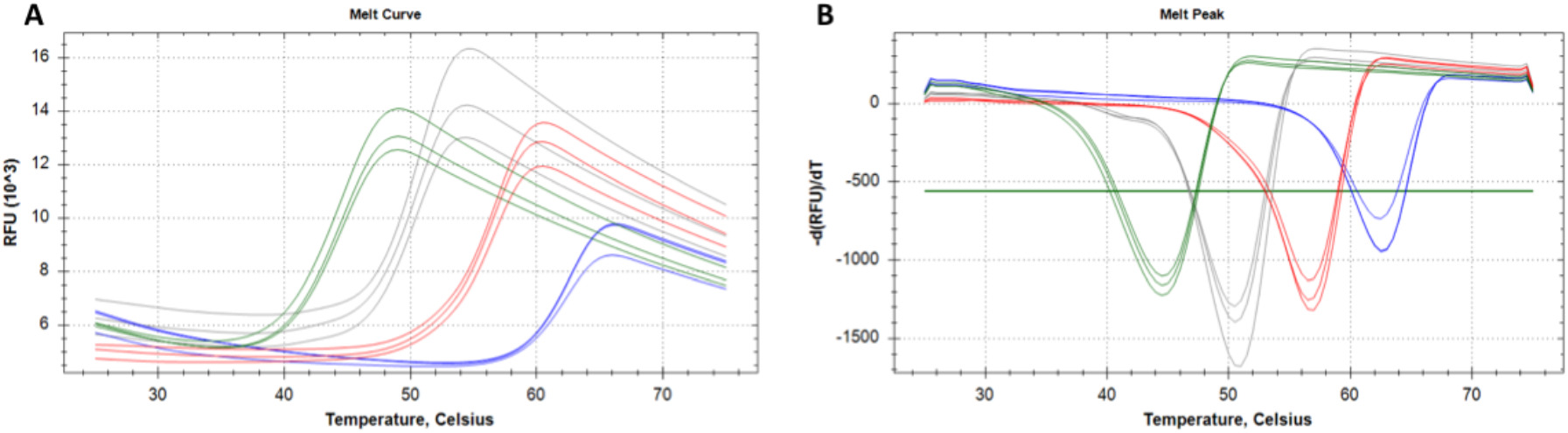
Biophysical characterization of MBZ binding to ABL1 *in vitro*. Representative unfolding curves (**A**) and positive derivative [d(RFU)/dT] curves (**B**) of fluorescence-based TSA performed with 4 μM of the kinase domain of ABL1 alone (*green*) or incubated with 200 μM of MBZ (*grey*), imatinib (*red*) or dasatinib (*blue*) over a temperature range of 10-95°C. *RFU*, relative fluorescence unit.

**Supplementary Figure 2.**
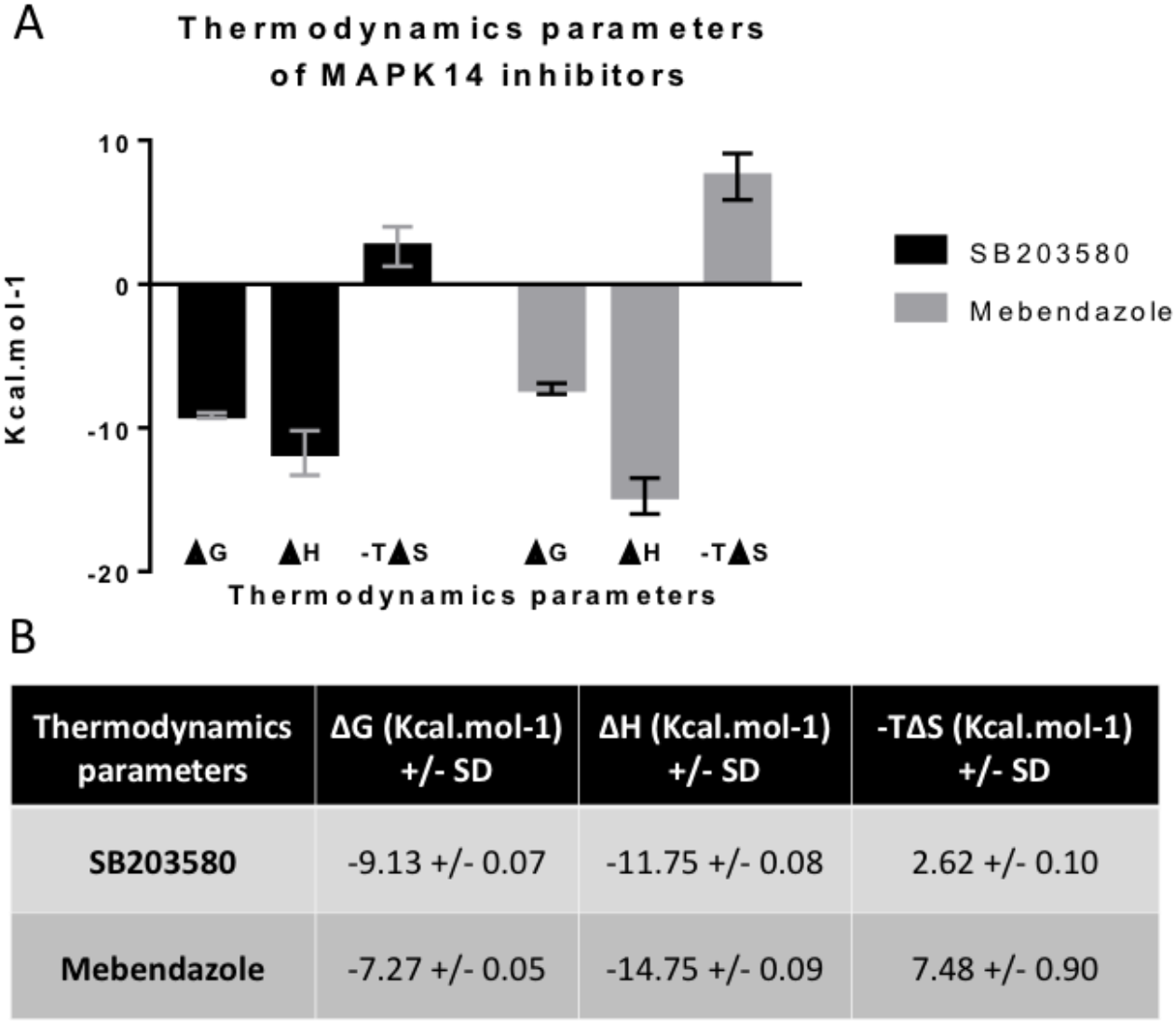
Thermodynamics parameters of MBZ binding to MAPK14. (**A**) Histogram of the thermodynamics parameters measured by ITC for SB203580 (*black*) and MBZ (*grey*) binding to MAPK14. (**B**) Summary table of the thermodynamics parameters (mean of at least 3 independent experiments).

**Supplementary Figure 3.**
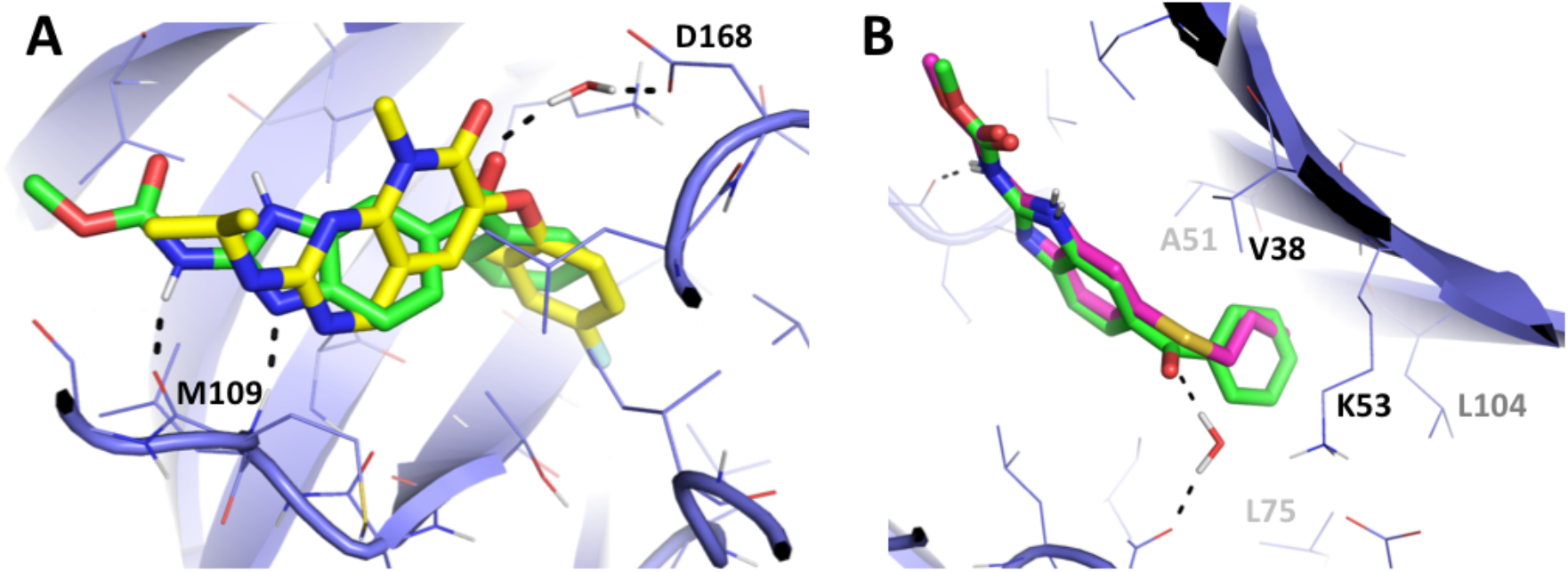
Binding mode comparisons of MBZ with 3FLY ligand and albendazole. (**A**) Comparison of the binding mode of MBZ (*green*) with the pyrido-pyrimidin inhibitor (*yellow*) from the 3FLY crystal structure. (**B**) Comparison of the binding mode of MBZ (*green*) and albendazole (*pink*). Ligands and surrounding protein sidechains are displayed as sticks and lines, respectively. The backbone from the protein is shown as cartoon representation.

**Supplementary Figure 4.**
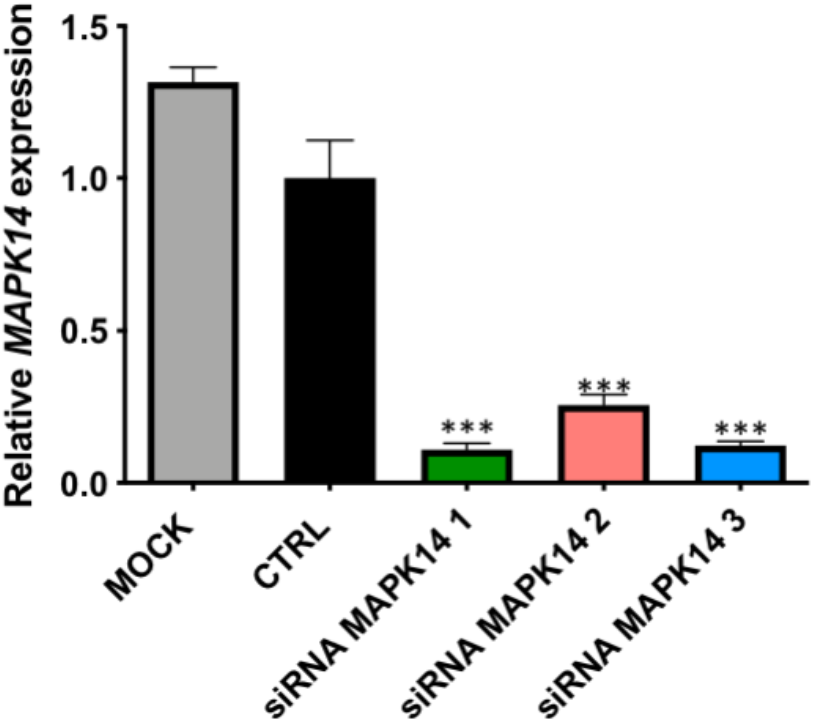
*MAPK14* gene silencing by RNA interference. Histogram of *MAPK14* gene expression, relative to housekeeping gene *YWHAZ*, in DsRed-expressing U87 cells following 48h transfection with negative ctrl (CTRL; *black*) and 3 different MAPK14 siRNA sequences (untransfected, MOCK; *grey*). Mean of 3 independent experiments +/- s.d; ***, p<0.001.

**Supplementary Table 1.** Average gene expression of MBZ putative targets in glioblastoma and normal brain tissue.

| Gene name | Average gene expression<br>normal tissues | Average gene expression<br>Glioblastoma subtypes | Fold<br>change | Two-way<br>Anova |
| --- | --- | --- | --- | --- |
| BLM | 1.22 | 6.25 | 5.1 | < 0.0001 |
| TP53 | 2.15 | 6.19 | 2.9 | < 0.0001 |
| KDR | 1.98 | 5.11 | 2.6 | < 0.0001 |
| NFKB1 | 2.36 | 5.92 | 2.5 | < 0.0001 |
| GMNN | 3.36 | 8.11 | 2.4 | < 0.0001 |
| CREBBP | 2.98 | 7.15 | 2.4 | < 0.0001 |
| HIF1A | 4.57 | 10.66 | 2.3 | < 0.0001 |
| NPC1 | 3.21 | 7.45 | 2.3 | < 0.0001 |
| MAPK14 | 3.34 | 6.84 | 2.1 | < 0.0001 |
| LMNA | 3.5 | 6.75 | 1.9 | < 0.0001 |
| ABL1 | 4.23 | 7.45 | 1.8 | < 0.0001 |
| MAPK1 | 5.32 | 7.82 | 1.5 | < 0.0001 |
| TEK | 3.18 | 4.43 | 1.4 | 0.0176 |
| HTT | 4.10 | 5.10 | 1.2 | 0.0230 |
| LGR3 | 0.29 | 0.51 | 1.8 | 0.9999 |
| FRAP1 | 3.53 | 3.94 | 1.1 | 0.9507 |
| ALDH1A1 | 5.12 | 4.96 | 1.0 | 0.9995 |
| ADORA2A | 4.24 | 3.26 | 0.8 | 0.1587 |
| THPO | 1.62 | 0.98 | 0.6 | 0.2326 |
| NPSR1 | 0.02 | nd |  |  |
| SLC6A3 | nd | 2.1 |  |  |

**Supplementary Table 2.** Pearson correlation between the cytotoxic activity of benzimidazole agents and their inhibitory effects on ABL1, ERK2 and MAPK14 in 4 GBM cell lines.

| Cell line | Pearson correlation ($R^2$ ; p value) | | |
| --- | --- | --- | --- |
|  | ABL1 | ERK2 | MAPK14 |
| U87 | **0,488**<br>p=0.30 | **0,009**<br>p=0.91 | **0,746**<br>p=0.14 |
| U87vIII | **0,810**<br>p=0.10 | **0,358**<br>p=0.40 | **0,996**<br>p=0.002** |
| T98G | **0,874**<br>p=0.065 | **0,518**<br>p=0,28 | **0,966**<br>p=0.017* |
| U251 | **0,611**<br>p=0.22 | **0,032**<br>p=0.82 | **0,821**<br>p=0.094 |

